# Finding the P3 in the P600: Decoding shared neural mechanisms of responses to syntactic violations and oddball target detection

**DOI:** 10.1101/449322

**Authors:** Jona Sassenhagen, Christian J. Fiebach

**Affiliations:** Department of Psychology, Goethe University Frankfurt, Frankfurt am Main, Germany; Brain Imaging Center, Goethe University Frankfurt, Frankfurst am Main, Germany

**Keywords:** ERP, multivariate pattern analysis (MVPA), language, syntax, P3, P600, domain specificity

## Abstract

The P600 Event-Related Brain Potential, elicited by syntactic violations in sentences, is generally interpreted as indicating language-specific structural/combinatorial processing, with far-reaching implications for models of language. P600 effects are also often taken as evidence for language-like grammars in non-linguistic domains like music or arithmetic. An alternative account, however, interprets the P600 as a P3, a domain-general brain response to salience. Using time-generalized multivariate pattern analysis, we demonstrate that P3 EEG patterns, elicited in a visual Oddball experiment, account for the P600 effect elicited in a syntactic violation experiment: P3 pattern-trained MVPA can classify P600 trials just as well as P600-trained ones. A second study replicates and generalizes this finding, and demonstrates its specificity by comparing it to face- and semantic mismatch-associated EEG responses. These results indicate that P3 and P600 share neural patterns to a substantial degree, calling into question the interpretation of P600 as a language-specific brain response and instead strengthening its association with the P3. More generally, our data indicate that observing P600-like brain responses provides no direct evidence for the presence of language-like grammars, in language or elsewhere.

## Introduction

Among the first discovered brain signatures of language processing was the influential observation of a positive-going deflection of the event-related brain potential (ERP), ~600 msec following syntactic violations, in a seminal study by Osterhout and Holcomb (1992; see also Hagoort, Brown, & Groothusen, 1993). This ERP component is now referred to as the P600 (for a review, see Kuperberg, 2007) and is associated with cognitive processes such as structural reanalysis, repair, or integration following syntactic violations or ambiguities (e.g., Friederici, 2011; Molinaro, Barber, & Carreiras, 2011; Brouwer, Fitz, & Hoeks, 2012). It is central to influential models of language processing (e.g., Friederici, 2002) which assume sequential processing stages, each realizing cognitive operations in one (more or less modular) linguistic domain.

The P600 is also reported for other modalities. By *reverse inference* (Poldrack, 2006; Kappenman & Luck, 2012), P600s are routinely taken as evidence that non-language domains rely on language-like syntax, or even involve the same neurocognitive modules supporting language processing. For example, Cohn, Paczynski, Jackendoff, Holcomb, and Kuperberg (2012) report a P600 for structurally malformed cartoon sequences, concluding that “the brain engages similar neurocognitive mechanisms to build structure across multiple domains” (p. 63). Patel (2003), investigating out-of-key chords in music, concludes that “the P600 reflected processes of structural integration shared by language and music” (p. 5). Christiansen, Conway, and Onnis (2012) interpret a P600 during nonlinguistic sequential learning as indicating that “the same neural mechanisms may be recruited for both syntactic processing of linguistic stimuli and sequential learning of structured sequence patterns more generally” (p. 1). Núñez-Peña and Honrubia-Serrano (2004) conclude from a P600 following arithmetic rule violations that “similar neurophysiological process could be required for the processing of violations in numerical sequences and in linguistic syntactic structures” (p. 130). We do not necessarily doubt such claims; it is possible that similar processes are taxed by both linguistic and nonlinguistic structural processing. However, we argue that evidence from the P600 may be insufficient to support such claims.

### The P600 as P3 Hypothesis

The P600 is hardly distinguishable from EEG patterns found in tasks with no requirement – or even possibility – for language-like structural processing. It particularly resembles the P300 ERP (Sutton, Braren, Zubin, & John, 1965; Luck, 2005) – specifically a late subcomponent called P3b (e.g., Debener et al., 2005; henceforth: P3) – which is prototypically observed in “oddball” paradigms requiring explicit detection of rare target stimuli, regardless of sensory modality. P600 and P3 both are characterized by long latencies (>300 msec) and a topographical distribution over centro-parietal scalp electrodes. Both ERP components share functional properties: The P3 follows salient (e.g., surprising or task-relevant) events (Nieuwenhuis et al., 2005) – which syntactic violations without doubt are – and both effects are attenuated for stimuli that are not task-critical (e.g., Osterhout, McKinnon, Bersick, & Corey, 1996; Hahne and Friederici, 2002; Osterhout, Allen, Mclaughlin, & Inoue 2002). The P600 is typically observed at later latencies, but this difference in latency is usually overstated – the P600 often peaks around 600 msec (e.g., in Osterhout & Holcomb, 1992), the P3 often around 500 msec (e.g., Makeig et al., 2004). Also, some very long-latency P3s – of 1 s or more – have been reported (e.g., O’Connell et al., 2012), and are, most importantly, expected for complex tasks, because the P3 latency is highly variable and depends directly on task complexity. The P3 was first observed – at ~300 msec – for a simple light flash experiment (Sutton et al., 1965). Any task more complex than that will induce a later P3 latency (as discussed by Verleger et al., 2005). Given that syntactic processing is substantially more complex than the detection of light flashes, differences in time course between P3 and P600 are thus also not a principled argument against shared underlying cognitive or neural processes. This dependence of P3 latency on task demands holds also on a single-trial level: if an overt task is given, P3 latency is correlated with response time (Verleger et al., 2005). The same holds for the latency of the P600 (Sassenhagen, Schlesewsky, Bornkessel-Schlesewsky, 2014; Sassenhagen & Bornkessel-Schlesewsky, 2015).

Further, just as the P3 is absent for nondetected targets (Nieuwenhuis et al., 2005), there is no P600 elicited for not-consciously detected syntactic violations (Batterink & Neville, 2013; Hasting & Kotz, 2008) or if linguistic errors can be expected due to, e.g., a non-native accent (Hanulíkova, van Alphen, van Goch, & Weber, 2012). The P600 is also elicited by sufficiently salient semantic violations (e.g., Münte, Heinze, Matzke, Wieringa, & Johannes, 1998; Kolk, Chwilla, van Herten,& Oor, 2003; Faustmann, Murdoch, Finnigan, & Copland, 2005; Szewczyk & Schriefers, 2011) and by a much-discussed class of phenomena on the edge of semantics and syntax called “reversal anomalies” (e.g., Kim & Osterhout, 2005; Kuperberg, 2007). Combined, these data suggest that subjective salience is *the* critical determinant for eliciting the P600. This account predicts that by modifying the subjective salience of a *semantic* manipulation, a P600 can be induced in semantic contexts. This is indeed the case; when semantic violations are made salient by, e.g., making them more explicit (van de Meerendonk, Kolk, Vissers & Chwilla, 2010) or task relevant (Kuperberg, 2007), a P600 is induced for semantic manipulations in sentence processing contexts.

Some authors (including Münte et al., 1998; Coulson, King, & Kutas, 1998; van de Meerendonk, Kolk, Vissers, & Chwilla, 2010; Bornkessel-Schlesewsky et al., 2011) speculate that P600 and P3 reflect the same neurocognitive process, with longer latencies in the linguistic domain due to higher stimulus complexity. Other P600 researchers (e.g. Osterhout and Hagoort, 1999; Frisch, Kotz, von Cramon, & Friederici, 2003) reject such reductive accounts, arguing that available evidence indicates a genuine P600 component specific to, and consequently indicative of, language and/or structural processing. This debate is complicated by the difficulty of establishing difference vs. identity of scalp topographic ERP distributions using classical statistical approaches (Luck, 2005). Thus, the above-described practice of reverse inference persists and neurocognitive models of language have not embraced a domain-general interpretation of P600 effects. To resolve this debate, new methods of EEG analysis are needed to clarify whether P600 and P3 rely on similar neurocognitive processes.

### Multivariate decoding for testing shared neural patterns

We propose that Multivariate Pattern Analysis (MVPA; King & Dehaene, 2014) is one such method. MVPA algorithms learn to predict the experimental condition to which trials belong by reading out (*decoding)* the information content of multivariate neural activation patterns representative of these conditions. If MVPA classifiers trained on such patterns in one experimental contrast (e.g., eliciting a P3) achieve high accuracy when classifying trials belonging to a different contrast (e.g., eliciting a P600), this indicates substantive similarity of (at least some of) the underlying neurocognitive events (Kaplan, Man, & Greening, 2015; Grootswagers, Wardle, & Carlson, 2017). Unlike null hypothesis testing of (dis-)similarity of trial-averaged ERP patterns, MVPA, thus, directly tracks to which degree multiple neurocognitive events share neural patterns – over space and/or time and as MVPA analyses involve classifying every single trial, at the level of the individual trial. Furthermore, unlike ERP or distributional similarity-based analyses, MVPA provides a direct, positive quantification (i.e., an intuitively interpretable effect size) of pattern similarity – allowing estimation and null-hypothesis significance testing of the degree of pattern sharing. By generalizing this analysis approach over time, MVPA can test whether specific neural patterns, identified at one time point, *persist* or *reoccur* across time, revealing shared patterns even if time courses differ. This makes it possible to explore whether the same neural pattern occurs at different time points in two different experimental settings – as would be the case if P3 and P600 indeed reflect the same underlying cognitive or neural process. In doing this, MVPA does not rely on identifying any single underlying generator, or a combination thereof, but instead considers the aggregate scalp pattern, i.e., potentially also a combination of multiple underlying generators. We think that for examining the P600-as-P3 hypothesis, this is particularly important as both P3 and P600 effects are assumed to consist of a mix of underlying neural generator components (e.g., Makeig et al., 2004; Friederici, 2011).

Here, we use time-generalized multivariate EEG decoding to test if P3 patterns elicited in an Oddball experiment generalize to a typical P600 experiment. We provide statistically reliable evidence that P600 contrasts are to a substantial degree shared with P3 patterns, corroborating the P600-as-P3 hypothesis. In a second study, we replicate this finding and validate the specificity of the method by demonstrating that cross-decoding between P600 and P3 is much stronger than cross-decoding between P600 and EEG correlates of processing two other categories of stimuli known to reliably induce ERP effects (i.e., faces, or semantically incongruent words).

## Study 1

### Materials & Methods

Hypotheses, analysis protocol, and an example analysis of preliminary data were pre-registered prior to data acquisition with the Open Science Foundation at https://osf.io/j2efc/register/565fb3678c5e4a66b5582f67. The preregistration for Study 1 included only the generalization across time decoding (as the only outcome measure); all other measures – simplified presentations of the results already contained in the generalization across time decoding – were selected in response to comments on an earlier version of this manuscript. (Study 2, with results fully in accordance with Study 1 – see below – included in its pre-registration protocol this simplified analysis, replicating the results of Study 1.) The experiment was approved by the local ethics board.

#### Participants

27 students at Goethe University Frankfurt were measured after giving informed consent. Two exclusions for technical failures and low performance left n = 25 participants (6 male), all native speakers of German, neurologically healthy, 18 – 29 (median: 22) years of age, and right-handed (> 24/28 on a test for right-handedness based on Oldfield, 1971). Participants were recruited on-line and compensated via course credit or financially.

#### Experimental procedures

Participants took part in two consecutive experiments, i.e., a language experiment, modeled after Osterhout and Mobley (1995), to elicit a P600, and a visual Oddball detection experiment to elicit a P3, free of syntax or semantics. The P3 experiment was conducted after the P600 experiment to prevent biasing how the language experiment was processed. Stimulus presentation was controlled with the open source Python software PsychoPy (Version 1.18.2; Peirce, 2007; http://psychopy.org). Experiments were preceded by printed instruction forms and a short sequence of training trials.

*Stimulus materials: Language experiment.* The language experiment (Fig. 1A) compared German sentences with subject-verb agreement violations to grammatical control sentences (as in Osterhout and Mobley, 1995; see also e.g. Wassenaar et al., 2004). Both types of sentences were constructed using a simple Markov grammar, made available at https://osf.io/j2efc/register/565fb3678c5e4a66b5582f67. For each participant, for each condition, 72 sentences were randomly created, with a length between 3 and 9 words or noun phrases. For ungrammatical conditions, sentences were generated by switching the number of the verb (i.e., singular to plural or vice versa) so that it no longer agreed with the subject (see (1) and (2) for examples and translations). Sentences were otherwise syntactically well formed. Sentence (1) becomes ungrammatical at the position of the verb, but (2) at the position of the subject; the critical word around which EEG activity was analyzed is indicated in bold. In the stimulus material, the critical item appeared at the final position 15% of the time.

1. Glaube mir ruhig, dass die Anwälte lange (**warteten/*wartete**). Believe me that the lawyers long (**had/*has**) waited. (literal translation: ‘Believe me that the lawyers ha[**d/s**] waited for a long time.’)
2. Angeblich (schläft/*schlafen) **der Gauner**. Allegedly (sleeps/*sleep) **the criminal**. (literal translation: ‘According to rumors, the criminal sleep[**s**].’)

144 additional, heterogenous sentences (i.e., the Potsdam Sentence Corpus; Kliegl et al., 2004) additionally were presented as fillers. Sentences were presented in white font (approximately 1° high) on grey background via rapid serial visual presentation, 300 msec for single words and 400 msec for noun phrases (with noun and determiner on one screen). Words/phrases were separated by 300 msec of blank screen. 800 msec after the sentence-final word, a question mark was presented until subjects responded with a button press (left vs. right index finger). As in Osterhout and Mobley (1995, Exp. 2), participants were instructed to judge whether or not the preceding sentence was anomalous and/or inacceptable, but were not explicitly instructed to judge ungrammaticality or to look out for subject/verb agreement mismatches. 1 sec after the response, the next sentence was presented. Before the main experiment, five training sentences were presented. The 288 sentences (2 * 72 plus 144 filler sentences) were presented with a break after every 25 sentences, which lasted until participants initiated the next block via button press.

*Oddball experiment.* The Oddball experiment (Fig. 1B) also consisted of 288 trials, each a sequence of visually presented crosses (‘+’; presentation time 200 msec) varying in color, inter-stimulus interval 350 msec. While the experiment resembles typical Oddball paradigms (Squires et al., 1975), stimuli were grouped into distinct sequences, to increase similarity to the language experiment. The number of crosses per trial – randomly chosen to range from three to nine – approximately corresponded to the number of words or noun phrases (i.e., screens) in the sentences of the language experiment. On 50% of trials, one cross was randomly selected to be of the target color. When considering all crosses presented across all trials, 8.7% of items were targets, realizing the Oddball manipulation. As before, 800 msec after each sequence, a question mark was presented until participants indicated via button press if the designated target color had appeared in the preceding sequence. Following the response, the next sequence followed. The experiment involved three colors (red, yellow, and blue). Before each block of 25 sequences, an instruction was visually presented that defined the rare, i.e., target color for the next sequences. Another color was used as a non-target distractor (novelty condition; not analysed here), that was presented with equal frequency as the target trials. All other crosses in a block were of the remaining, common/standard color.

**Figure 1.**
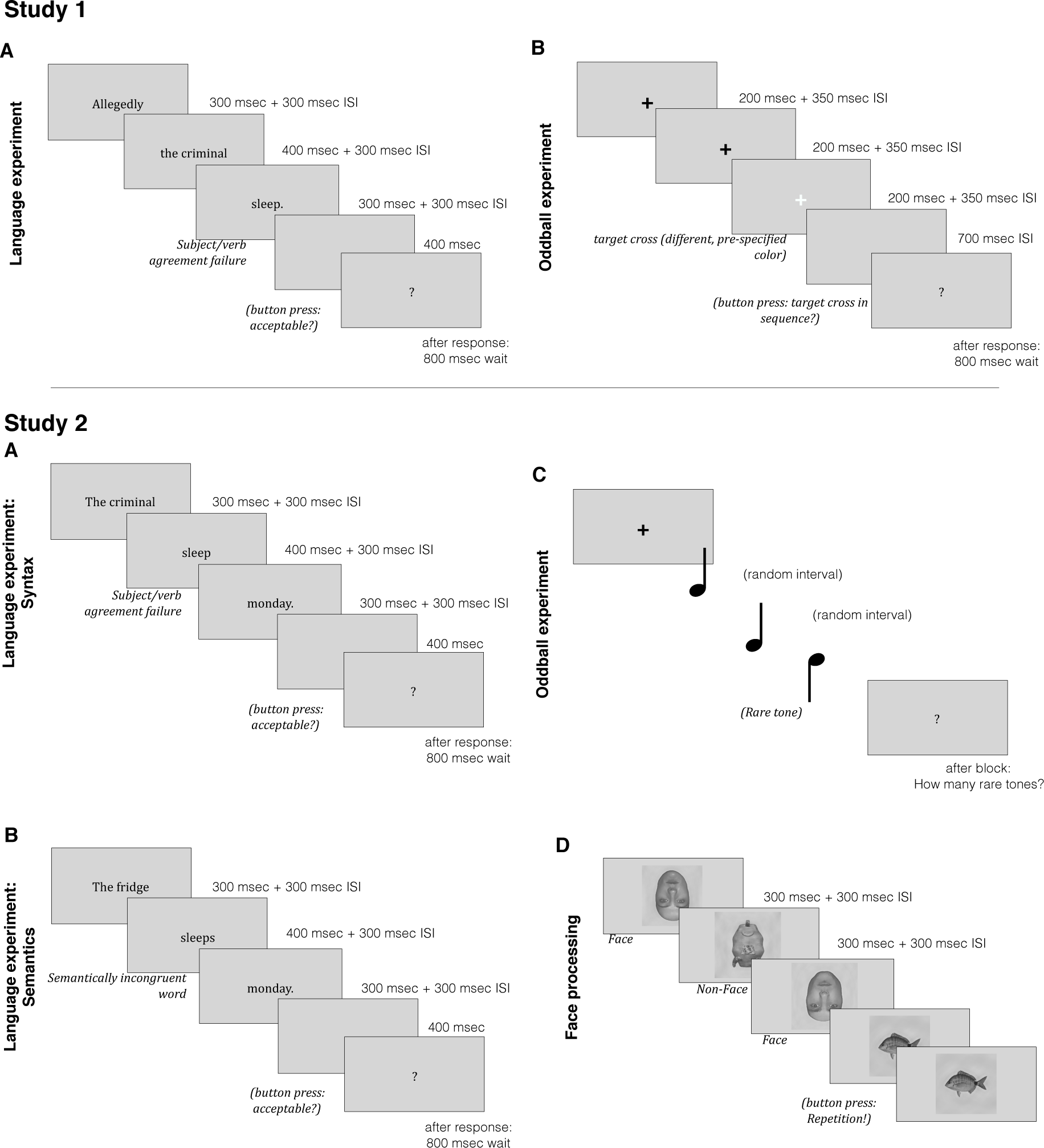
Experimental paradigms. *Top:* Study 1. **A**: Trial timing of sentence presentation in the syntactic violation experiment. While the example sentence indicates the general nature of the stimuli, the actual – German – stimulus sentences were heterogeneous with respect to structure and length; see online code repository for the full stimulus material. **B**: Outline of a trial from the Oddball experiment. Black crosses represent the standard color; target crosses (here: white) were of a different colour and could occur with equal probability at any sequence position but the first. ISI, inter-stimulus interval. *Bottom:* Study 2. Outline of the trial sequences of experimental tasks: the two language conditions (A, B), and the tone Oddball paradigm (**A**) and face processing task (**B**).

#### EEG data acquisition and analysis

EEG was recorded with 64 active Ag/AgCl electrodes (actiCAP; Brainproducts, Munich, Germany) positioned according to a 10/10 layout at 1,000 Hz sampling rate (BrainAmp DC amplifier; Brainproducts, Munich, Germany). Data were processed, following the initial pre-registration protocol, with MNE-Python (Gramfort et al., 2013; http://www.martinos.org/mne/) in iPython (Pérez and Granger, 2007). Data were cleaned of stereotypical artefacts via ICA (Jung et al., 2000; semi-automatic component selection via Corrmap, Viola et al., 2007), bandpass filtered between .1 and 30 Hz, downsampled to 200 Hz, re-referenced to linked mastoids, epoched around critical items; Oddball experiment: onsets of rare and common, but not novel crosses; language experiment: words at which the agreement mismatch became detectable vs. words at the same position in well-formed and congruent control sentences. Epochs were baseline corrected by subtracting the mean over a 500 msec pre-stimulus window, and then cropped to a 300 msec baseline for further analysis. Epochs with extreme amplitudes (> 150 uV) were dropped. As in the P600 reference study (Osterhout and Mobley, 1995), all remaining trials were entered into the analysis independent of behavioral response correctness. Epoch lengths were chosen according to the expected effects, i.e., 1 s (syntax) and 600 msec (oddball).

#### Multivariate Pattern Analysis

MVPA decoding with generalization across time was conducted similar to King and Dehaene (2014) and King et al. (2014). See Fig. 2 for an overview of our analysis approach. For the Oddball data of each subject, five independent trial subsets were created, with a similar balance of items per condition (i.e., stratified 5-fold cross-validation). In each fold, four of the five subsets were used as training set to *train* classifiers. Within each subject’s training set, a Logistic Regression (implemented in scikit-learn; Pedregosa et al., 2011; http://scikit-learn.org) with balanced class weights (so that items of the rarer class, i.e., Oddballs, received higher weights) was trained to separate standard from Oddball trials based on EEG activity (Fig. 2A1). This was conducted independently at each time point during a temporal window of ±50 msec around 400 msec, i.e., the expected center of the P3 effect (Nieuwenhuis et al., 2005). Each electrode corresponds to one feature in these analyses, so that per subject, 20 classification analyses (i.e., 100 msec sampled at 200 Hz) were conducted based on 63 features (i.e., electrodes) each. The resulting P3-classifiers were then *tested* by letting them predict trial type - i.e., standard vs. Oddball - for each time point of each trial of the remaining fifth subset (the test set; Fig. 2A2), based on their pattern of EEG activity (Fig. 2A3). To quantify the accuracy of these predictions, across all trials of the test set, the probabilities that the P3-classifiers had assigned to the positive class were scored by computing the area under the curve of the receiver-operating characteristic (ROC AUC), a nonparametric measure robust to class imbalances. This was repeated five times (once per fold), so that each trial subset served as testing set once (see Fig. 2A2), and ROC AUC scores were averaged across the five folds. This resulted in one time series per participant, corresponding to the strength of differential EEG activity separating Oddball from standard trials, at each sample point of the trial.

The critical analysis was to evaluate whether these subject-specific P3-trained classifiers generalize to the respective participants’ syntactic P600, i.e., whether they could also categorize trials from the language experiment into grammatical violations vs. grammatically correct control sentences (see also Fig. 2C2). For this, we tested for each participant how well their P3-classifiers can predict, based on EEG activity elicited by the critical word, if it stems from a language trial with vs. without a syntactic violation, using all grammatical and ungrammatical trials. For each subject, the average of all five P3-classifier accuracies (resulting from the five-fold cross validation; see above) was used for classifying the trials from the sentence experiment. As described above, to learn the P3 patterns, classifiers were trained on the P3 effect and were then tested at each time point of each sentence trial. A prediction was counted as correct if a violation trial was classified by the P3-classifier as an Oddball, or a control sentence trial as standard; otherwise the prediction was counted as incorrect. Predictions were scored, as described before, via the averaged ROC AUCs. Above-chance decoding here indicates that a neural activation pattern observed during the P3 effect in the Oddball experiment is also reliably found in the language experiment. Below-chance decoding would indicate that EEG patterns elicited by the oddball condition are stronger in the control condition than in the violation condition.

As noted, the described cross-experiment decoding procedure yielded one time series per participant, corresponding to the degree to which P600 trials could be classified correctly by applying the P3-classifiers to EEG activity from the language experiment, at each sample point in the EEG epoch. For statistical evaluation, the mean performances of these classifiers during the P600 effect (at 700±50 msec) were compared across participants with a one-sided signed-rank test (because accuracy scores are not normally distributed). Following the P600-as-P3 hypothesis, we predicted that if the P600 shares neural processes with the P3, P3 classifiers should be able to classify syntactic violation trials.

**Figure 2:**
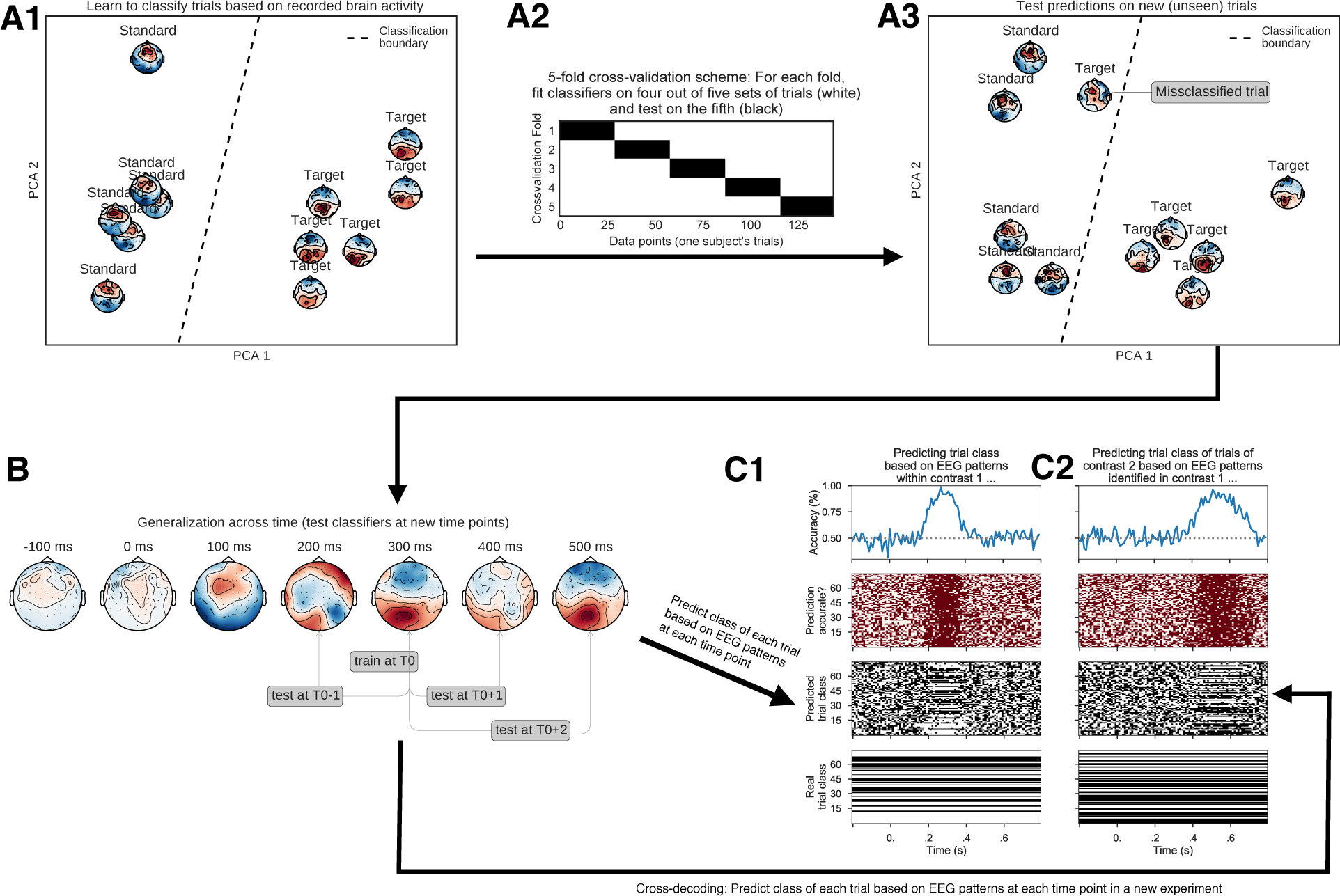
Demonstration of decoding procedure (simulated data). **A**: Cross-validated training and scoring of classifiers. (**A1**) Training of a machine learning pattern classification algorithm (classifier). Each topographical plot represents neural activity at one time point for one trial of the training set. Multivariate pattern classification considers the EEG pattern across all sensors. Classifiers identify the best-possible separation between trials of the two classes based on their patterns of brain activity. The separation boundary is represented by the dashed line. (**A2**) 5-fold cross-validation procedure: To avoid circularity, five independent classifiers are each trained on subsets of 80% of the data (training set; white) and validated on the remaining 20% of previously unseen trials (testing set; black). (**A3**) Scoring these predictors on the testing set results in an estimate of classification accuracy. Here, topographies represent the previously unseen trials of the testing set. Trials to the left of the separation boundary (which is learned during training; cf. **A1**) are predicted to belong to the Standard class; one trial has been misclassified. (**B**, **C1**) Generalization across time: Within each of the five folds, trained classifiers (trained at a specific point in time; e.g., at 300 msec in the example in B) are tested by predicting trial class membership of each test trial at each time point of the trial (**C1**, third row). These results are scored for accuracy (**C1**, second row) relative to the actual trial category (**C1**, bottom row), yielding a time series of classification accuracies (**C1**, top). This indicates in which time windows an EEG pattern is present (here: ~400-600 msec). (**C2**) In cross-experiment classification, the same approach can be used to make predictions based on brain activity from a different experiment or experimental contrast. The simulated example shows the occurrence of the identical pattern as in **C1**, however in a later time window.

In addition, as explained above, besides for time-resolved decoding, we also conducted temporal decoding with *generalization across time*. Each classifier (trained at one time point) was evaluated at every other time point. Here too, classifiers trained on Oddball data were evaluted on both Oddball and on language experiment data (resolving across time in both cases). The resulting matrices indicate at which time point the pattern identified in the Oddball data reoccurs in the testing data – i.e., not only for selected classifiers (such as the peak P3 classifier), but for all classifiers, at all time points. For example, an above-chance decoding score for Oddball classifiers trained at 320 msec and tested on the data from the language experiment at 700 msec indicates that EEG patterns contributing to the P300 reoccur during the P600 time window in syntactic violation trials. The results of this procedure were averaged across participants and plotted as heatmaps (see below). Following the original pre-registration, all time points following stimulus onset were tested for statistical significance with a rank-sum test against chance across participants. For statistical thresholding of cross-experiment decoding from Oddball P3 to syntactic violation P600, only those classifiers which were above chance in the within-experiment decoding of the P300 were included.

## Results

### Behavioral results

In accordance with the pre-registration, because the focus of the study was on well-known EEG effects and their relationships, behavioral results were not assessed in detail. However, one subject was excluded for having < 75% accuracy on the syntactic judgement task.

### Event-Related Potential (ERP) results

As in earlier studies (e.g., Nieuwenhuis et al., 2005), Oddball relative to standard trials in the P3 experiment elicited an ERP positivity with a centro-parietal distribution (Fig. 3A), appearing as early as 350 msec and peaking around 400 msec. Consistent with classical P600 ERP reports (e.g., Osterhout et al., 1992, 1995, 2002), the ERP contrast between syntactic violations and grammatical control sentences in the language experiment was characterized by a late positivity over central and parietal electrodes, beginning ~500 msec post onset of the ungrammatical target word (Fig. 3B). The P3 was stronger than the language effect (i.e., ~15 vs. ~6 uV at their respective peaks; the 95% confidence interval at the P3 effect peak excluded the upper range of the 95% confidence interval of the P600 effect peak).

### Time-Resolved Decoding results

The critical hypothesis test of this study involves the cross-decoding between experiments: Can classifiers trained on the P3 effect significantly predict the experimental condition (i.e., violation vs. control) when applied to language trials? P3-to-P600 decoding was better than chance, beginning at ~500 msec. Statistical evaluation in the pre-registered time window of 650-750 msec yielded a classification accuracy ~60% that was highly significant (p < .0001; Fig 3C).

In time-generalized decoding (within-P3 decoding; Fig 3D), the P3 appears as a broadly quadratic pattern, indicating a single effect persisting in time. This replicates the quadratic pattern observed already by, e.g., King et al. (2014) for the P300. Considering the between-experiment generalization (i.e., P3->P600 decoding), P3 classifiers trained from ~250 msec onwards could successfully decode syntactic violation trials based on data from ~450 msec onwards. The rectangular nature of the effect (p < .001) around Y = 400 msec and X = 750 msec (i.e., including the pre-specified time windows) indicates that any MVPA classifiers trained throughout the time window of the Oddball/P3 effect can successfully classify trials as syntactic violations vs. controls based on EEG activity throughout much of the P600 time window. This indicates that there is only one pattern occurring in the P3 contrast which also persists through all of the P600 contrast.

**Figure 3:**
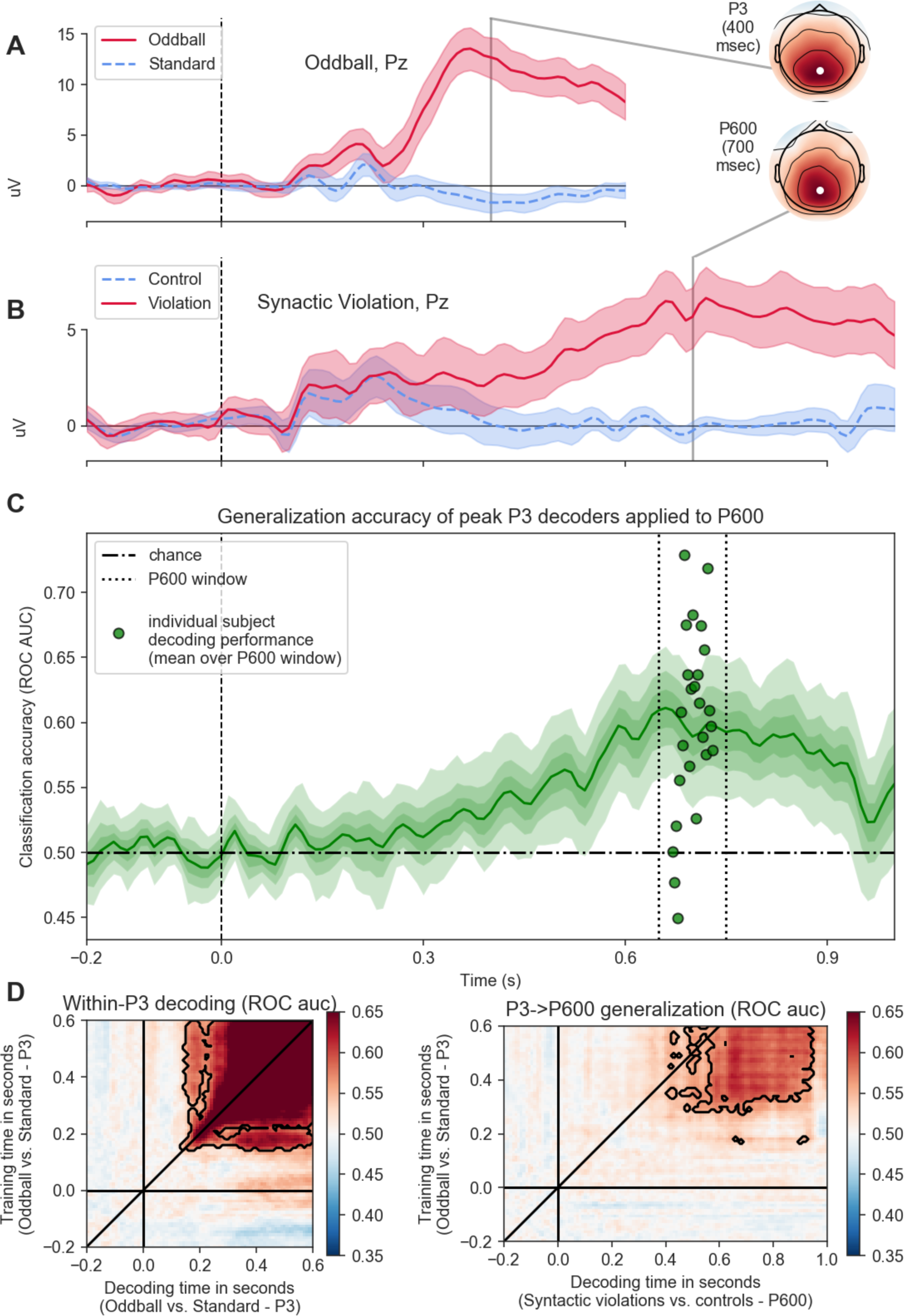
Results, Study 1. Top: ERPs for Oddball (**A**) and language (**B**) experiments, respectively, at a representative electrode (Pz). Interpolated scalp maps show the critical contrast at the pre-registered time windows used for statistical and decoding analyses of P3 and P600 (i.e., 400 ± 50 msec and 700 ± 50 msec, respectively). Note the different scaling of y-axes. (**C**) Time series of cross-experiment decoding performance when classifying trials from the language experiment (i.e., grammatical vs. syntactic violations) by the classifiers trained on the P3 effect. P3 classifiers were trained on the critical time window of the P3 contrast (350-450 msec) and then applied to all time points of the language trials (as illustrated in Figure 2). Dashed vertical lines represent the pre-registered time window of 650-750 msec used for statistical evaluation of cross-decoding. Error corridors show bootstrapped confidence intervals (95% for ERPs; 68%, 95%, and 99% for decoding). (**D**) Generalization across time (GAT) results. Left: training time (X-axis) by testing time (Y-axis) decoding scores for classifying Oddball vs. Standard trials based on classifiers trained on the Oddball contrast (i.e., within-P3 decoding). The Y axis shows where in the training set the pattern occurs. The X axis indicates at what time points in the trial this specific pattern can be decoded. Successful decoding along the diagonal demonstrates the occurrence of an effect, analogous to ERP results, whereas off-diagonal decoding indicates temporal generalization in the sense of persistence of re-occurrence of an effect. Black outlines indicate significant regions after FDR correction (p < .001). Right: same, but applying P3-trained classifiers to the EEG data from the syntactic violation/language experiment.

## Interim discussion

Using time-generalized multivariate pattern analysis, we demonstrate that patterns of EEG brain activity characteristic of the Oddball-P3 are also activated during processing ungrammatical sentences, in the time window of the well-established P600 ERP effect. This indicates substantial overlap between the two components at the level of neurocognitive processes (as proposed by the P600-as-P3 account; e.g., Coulson et al., 1998; Sassenhagen et al., 2014).

However, these conclusions heavily depend on the specificity of demonstrated cross-decoding for P600 and P3. Put differently, the support of our results for the P600-as-P3 hypothesis would be questionable if other EEG responses would cross-decode to the P600 in similar fashion. MVPA cross-decoding of electrophysiological brain responses is novel (King & Dehaene, 2014) and to the best of our knowledge, no empirical data exist concerning the specificity of the method. To investigate the specificity of our cross-decoding result, we pre-registered and conducted a second study comparing P3-to-P600 cross-decoding to cross-decoding of two other well-known, reliably-elicited ERP responses with clearly different antecedent conditions and spatial topographies, the N400 effect (elicited by semantic violations; Kutas & Hillyard, 1980) and the N170 (elicited by faces; Bentin, Allison, Puce, Perez, & McCarthy, 1996).

We also cannot exclude that the specific format of our visual Oddball experiment (with mini-sequences similar to rapid serial-visual sentence presentation) may have favored P3-to-P600 cross-decoding. To exclude this, we implemented in Study 2 a classical auditory Oddball experiment without a trial-like sequential component. Also, a different set of linguistic stimuli was chosen in the language task to avoid potential confounds from post-sentence wrap-up effects (due to sentence-final placement of critical items in 15% of the sentences in Study 1). Study 2, thus, further served to replicate the primary finding and establish its robustness in the sense of domain-generality and invariance against specific details of trial structure or stimuli.

## Study 2

### Materials & Methods

The study, including hypotheses, specific analysis protocols, and an example analysis script of preliminary data (specifying the parameters for the actual analyses), was pre-registered with the Open Science Foundation at https://osf.io/7e93a/.

#### Participants

35 neurologically healthy, right-handed native speakers of German were measured, after giving informed consent according to protocols approved by the local ethics committee (inclusion criteria as in Experiment 1). Five exclusions due to technical failures left 30 participants (14 male; 18 – 30 years; median age: 22), as pre-registered. Sample size was chosen to guarantee >95% power for our main hypotheses according to a resampling-based investigation of data from Study 1.

*Language experiment: Syntactic and semantic violations.* Example stimuli are given in (3-6). For the language experiment, violation sentences were constructed from an initial set of 320 meaningful and grammatically well-formed German sentences, by either exchanging a noun or verb between two selected sentences so that at the position of the switched word, the sentence became highly implausible (semantic violations; 3, 4), or by exchanging the verb or a pronominal subject of a sentence for a morphosyntactically ill-fitting one from another sentence (syntactic violations; 5, 6). As in Exp. 1, the syntactic manipulation consisted in agreement violations; however, they were of a slightly different type, in part to avoid sentence-final violations. Note that (5) becomes ungrammatical at the position of the verb, but (6) at the position of the subject. The critical word around which EEG activity was analyzed is indicated in bold.

(3) Der Storch erspäht den [**Frosch/*Duft**] aus der Luft. the stork sees the [**frog/smell**] from the sky
(4) Den Keks [**isst/*entkräftet**] der Junge voller Genuß. The cookie [**eats/disproves**] the boy with passion (literal translation: “The boy enjoys eating/disproving the cookie.”)
(5) Der Garten [**blüht/*blühen**] schon im Frühling. The garden [**blossom/*blossoms**] already in the spring
(6) Den Kuchen mögen [**sie/*er**] ganz besonders. The cake like [**them/he**] very much “They/he particularly like the cake.”

Two experimental lists were constructed, each with 80 morphosyntactic violations, 80 semantic violations, 160 matching control, and 10 additional filler sentences, so that each critical sentence appeared in one list as a violation and in the other as a control. For each participant, one of the two sentence lists was chosen randomly. Sentences were presented phrase by phrase (e.g., keeping determiners and nouns on one screen), with timings and task modalities as for the sentence processing experiment in Study 1.

*Oddball experiment.* The oddball experiment was modelled after Debener et al. (2005) and consisted of a sequence of two sine wave tones, one of which was randomly (per subject) chosen to be the standard tone, accounting for 95% of presentations, and the other to be the rare tone (randomly chosen for 5% of trials). Tones were played for 340 msec, with a randomly chosen inter-stimulus interval of 960-1360 msec. Participants were queried about the number of rare tones after each blocks of 80 tones. After each block, participants could proceed to the next block via button press.

*Face processing experiment.* Three kinds of grayscale picture stimuli were presented to each participant: 8 inverted faces (to induce an N170 component; see Goffaux et al., 2006, see Sadeh & Yovel, 2010), 8 face-like non-face objects (Vuong et al., 2017), and 24 other non-face pictures, each presented 20 times. Pictures were presented visually in blocks of 30 items, for 300 msec each, followed by an inter-stimulus interval of 700 msec. After each of 30 blocks, participants could proceed to the next block via button press whenever they felt so. Participants had been instructed to respond via button press if an image was an exact repetition of the preceding image.

#### EEG Data Preprocessing

Data were preprocessed as described for Study 1, except that for visualization of ERP patterns (but not for decoding analyses), face processing trials were re-referenced to average reference (following standard procedures). EEG was again epoched around critical items; i.e., onsets of rare and common tones (Oddball experiment); onsets of words at which the (syntactic or semantic) mismatch became detectable, and respective words in control sentences (language experiment); faces vs. non-face objects (face processing experiment). Epoch lengths were pre-registered in accordance with the lengths of expected effects, i.e., 1.2 s (Syntax), 800 msec (Semantics), 600 msec (Oddball), and 300 msec (Face).

#### Multivariate Pattern Analysis

Data were analyzed as before, with (cross-)decoding performed from classifiers trained on every task to the syntactic violation contrast (i.e., Syntax/P600->P600, Oddball/P3->P600, Semantics/N400->P600, and Faces/N170->P600). Statistical evaluation was performed as in Study 1 in time windows of ±50 msec around the respective component’s center (i.e., 400, 700, 400, and 170 msec for P3, P600, N400, and N170, respectively); time windows were selected and pre-registered based on the literature. (Note that results turned out to be robust to selecting different windows.) To account for the possibility of polarity effects (i.e., the possibility that patterns like the N400 might more resemble the syntactic control trials that have relatively more negative-going ERPs than syntactic violation trials), which would lead to below-chance decoding performance, below-chance scores were ‘inverted’ (1 - score). Note that while the N400 is negative in polarity, the N170, while nominally a “negativity”, is under common referencing schemes (i.e., the bimastoid and the average references employed in this study) positive at more sensors than negative (i.e., at central and frontal ones; see Bentin et al., 1996, Fig. 3C; or this manuscript, Fig. 3G).

#### Additional (not pre-registered) analyses

First, to assess the (dis-)similarity between P3 and P600, P3-to-P600 cross-decoding was compared to within-P600 decoding statistically in the pre-registered critical time window of 700 ± 50 msec. Second, we further explored the new control conditions, by (i) comparing N400-to-P600 and N170-to-P600 cross-decoding to chance, and by calculating a t-test of the across-subject grand means of the differences (with time points as observations) between the two decoding time series, for P3, N400, and N170, compared to P600-to-P600 decoding. To visualize the relative strength of decoding the P600 based on the other patterns, we furthermore calculated the ratio of decoding scores for language trial data for each decoder relative to the decoding scores for the language-data trained decoder. Third, as a control analysis, we inverted our decoding pipeline to investigate if cross-decoding from P600 to P3 was stronger than cross-decoding from P600 to N170 or N400, at the respective components’ centers. Finally, we calculated generalization across time matrices for Study 2 as for Study 1 (only shown in supplement).

## Results

**Fig. 4:**
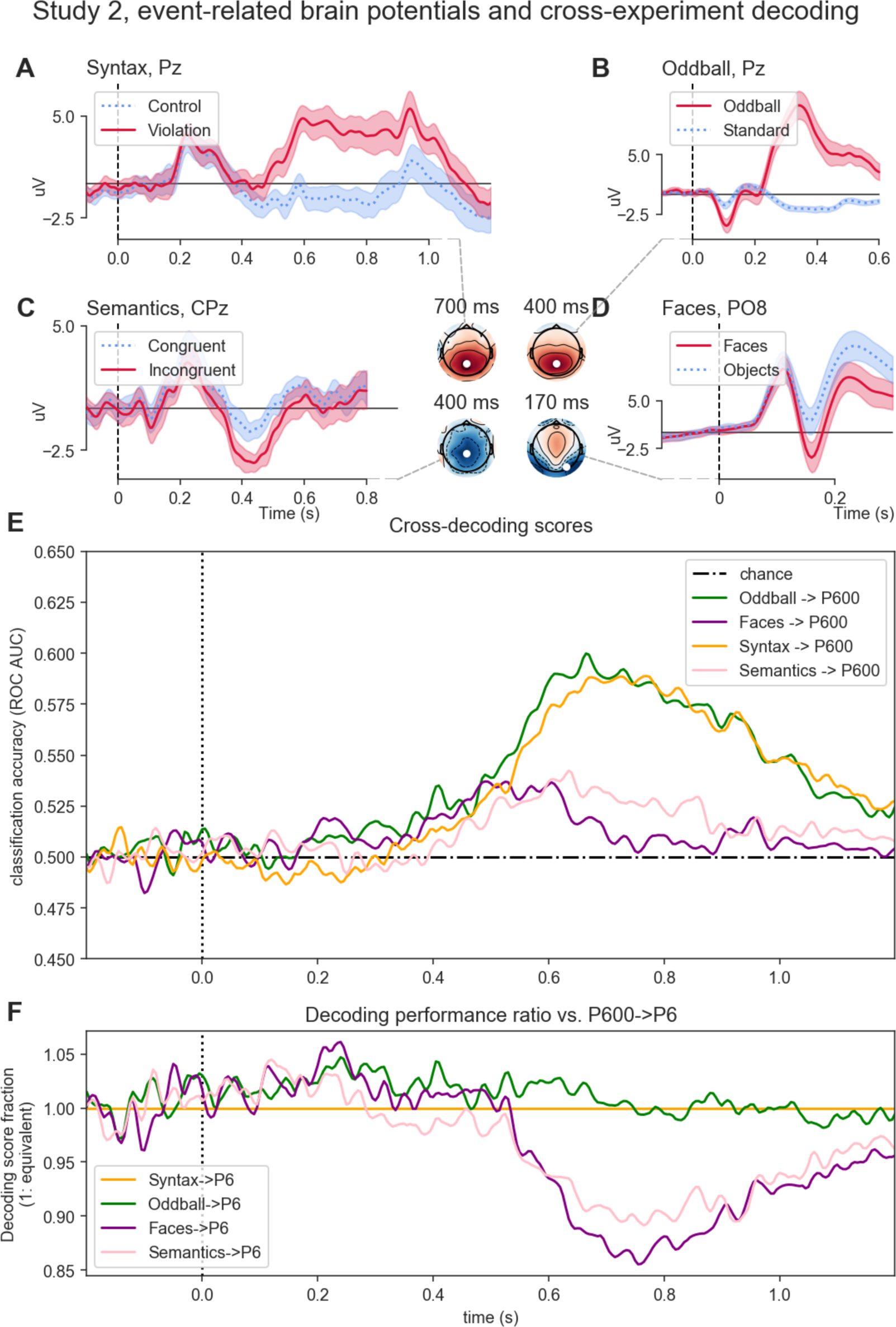
Results, Study 2. Top: ERPs for P600 (**A**), P3 (**B**), N400 (**C**), and N170 (**D**) contrasts, shown at representative electrodes. Interpolated scalp maps show the critical contrast at the pre-registered time windows used for statistical and decoding analyses. Note that the scaling of x- and y-axes differs between the plots. Error corridors show bootstrapped 95% confidence intervals for ERPs. (**E**) Decoding scores when classifying Syntactic violation vs. Control trials in the language experiment, based on classifiers trained on the peak effects in the four sub-experiments. (**F**) Same data as in (**E**), but, for each experiment, expressed as the ratio between P600-to-P600 decoding and decoding based on that experiment; i.e., how much worse are decoders trained in any one experiment when applied to P600 data, compared to decoders trained on P600 data itself.

### Behavioral results

As in Study 1, and in accordance with the pre-registration, no detailed analysis of behavioral results was performed. All subjects performed above chance accuracy, so no exclusion for low performance was necessary.

### Event-related brain potentials (ERPs)

Figure 4A-D displays the ERPs elicited by control vs. syntactic violation sentences, congruent sentences vs. semantic violations, standard vs. Oddball tone trials, and faces vs. non-face objects. ERPs to auditory Oddballs and syntactic violations are highly comparable to those observed in Study 1. Semantic violations induced a centro-parietal negativity peaking slightly after 400 msec (Figure 4C). Faces vs. non-faces induced the expected differential ERP patterns in the form of a N170 (Figure 4D). As already mentioned above, the scalp distribution of the N170 involves broad regions of positive-going potential differences (with 63% of electrodes positive).

### Multivariate Pattern Analysis

Figure 4E shows time courses for the accuracies of decoding syntactic violations, based on classifiers trained on every critical experimental effect. This plot, thus, shows to what extent, at any given time point, EEG patterns typical for the P600 violation effect itself (green), the oddball P3 (orange), semantic violations (pink), or for a face-specific N170 (purple) can classify whether a given EEG signal was elicited by the target word in a grammatically correct control sentence or a syntactic violation trial.

Decoding within the syntax experiment showed peak decoding accuracies >60% at ~700 msec (median ROC AUC: >.58, p < .0001). This within-P600 decoding can be understood as defining an approximate upper bound for how well syntactic violation occurrences can be predicted from corresponding brain patterns. Classifying syntactic violation vs. control trials based on the auditory oddball P3-classifier also showed peak decoding accuracies ~60% at ~700 msec (median: .6, p < .0001), with a near-identical time course (r > .98; 95% CI: 0.958, .99). This indicates that P3->P600 decoding reaches the upper bound for P600 decoding observed in P600->P600 decoding. As predicted, P3-to-P600 cross-decoding was substantially stronger than the accuracies of both N170-to-P600 and N400-to-P600 cross-decoding (Oddball vs. Face: p < .001; Oddball vs. semantic violation: p < .0001; tested in the pre-registered time windows of interest; cf. Figs. 4E, F).

### Non-Pre-Registered Analyses

The first additional (i.e., not pre-registered) analysis tested for differences in decoding accuracies between within-experiment decoding of syntactic violations (P600-to-P600) and P3-to-P600 cross-decoding, but did not observe any significant differences (p > .24; tested in pre-registered P600 time window). Note that this analysis is very similar, but not identical to the correlation analysis of the decoding time courses reported in the preceding paragraph. Second, we investigated in slightly more detail the cross-decoding from N400 and N170, beyond the co-registered test: Albeit significantly weaker than P3-to-P600 cross-decoding (see previous section), cross-decoding from Face and Semantic violation trials to P600 (Fig. 4F, pink and purple lines) was significantly different from chance (Faces: p < .028, Semantics: p < .0058; P600 time window). When statistically comparing time courses rather than pre-selected time-windows, P600-to-P600 decoding significantly differed from of N170-to-P600 and N400-to-P600 cross-decoding (both *t*(29) > 2, p < .05), but not from P3-to-P600 decoding time courses (|*t*|(29) < 1, p > .05). Finally, examining cross-decoding from P600 to all other experimental contrasts (i.e., testing a P600-trained classifiers in the respective time windows of interest for each ERP effect), we observed strong (>65%; p > 0.001) cross-decoding from P600 to P3, again significantly (p < .0001) stronger than when cross-decoding from P600 to either N400 or N170; see supplement, Fig. 1. Generalization-across-time MVPA analysis for Study 2 are shown in the supplement, Fig. 2.

## Discussion

The starting point for the present study was the long-standing debate about the relationship between P600 – particularly that observed in (morpho-)syntactic violations (e.g. Osterhout and Mobley, 1995; Osterhout and Holcomb, 1992) – and the P3 ERP component (cf. Coulson et al., 1998; Sassenhagen et al., 2014). We applied time-generalized MVPA decoding across experimental tasks, in the same participants, and demonstrate in two independent datasets that EEG patterns elicited by rare stimuli in Oddball experiments account for the neurophysiological signals elicited by syntactic violations in sentences, with near-identical time courses as within-P600 decoding. Phrased differently, decoding P600 trials (i.e., with respect to their nature as control vs. syntactic violation trials) by classifiers trained on P3 data was just as good as decoding P600 trials based on P600 data itself, with nearly indistinguishable performances and decoding time courses.

Across the two experiments, we showed this for visual and for auditory Oddballs, with multiple different kinds of syntactic violation sentences. Combined, these results indicate that the P600 shares to a substantial degree neurocognitive processes with the domain-general P3 elicited during conscious detection of salient events. Face- and semantic violation-induced EEG patterns, in contrast, did not allow comparable cross-decoding, indicating that the P600 shares neural patterns specifically with the P3 ERP effect. Note that the N170 component, which is, despite its name, positive at more sensors than negative (in our case: 63% positive), did not afford substantially better cross-decoding than the N400 (0% sensors with positive ERP differences).

This result may come at no surprise for some readers. Descriptively, P3 and P600 often have very similar topographies (as already noted by, e.g., Osterhout et al., 1992); and N400 and P600, and even N170 and P600, have visually distinct topographies. However, we want to emphasize the asymmetry of the test employed here: The P600-as-P3 perspective (in its many iterations; including, e.g., Coulson et al., 1998; Kolk et al., 2003) predicts that for any person, their P600 will correspond strictly to their P3 (instead of, e.g., the grand-averaged components looking similar); these theories would have been falsified by any other outcome to this study than the one we observed. In contrast, other theories make no such predictions. Secondly, MVPA, in particular time-resolved MVPA cross-decoding, can deal with the case that effects characteristic of two experimental contrasts show up in different time windows. MVPA, thus, provides a principled, explicit measurement for quantifying ‘similarity’; in fact, so we argue, MVPA should be the standard measurement for this research question. Older techniques – such as correlations between scalp maps, or (non-significant) difference tests, cannot address this question in a principled, statistically adequate manner. In contrast, cross-decoding provides scientific and accurate error bounds on how strongly two EEG patterns resemble each other. In our case, the results of the MVPA decoding analysis argue for strict identity between at least the peaks of the P600 and P3. By this, our results also provide more insight into the functional nature of the P600 than could be obtained from a source-level analysis, e.g., from an fMRI investigation of syntactic violations. For this reason, we on purpose did not approach the P600-as-P3 hypothesis by searching for commonalities in their neuroanatomical localizations; instead, we aimed to compare two ERP components – to test the theory that P600 and P3 share a neurocognitive substrate.

While our between-experiment decoding results do not *prove* the functional identity of the two components, it is a strong *test* of the identity hypothesis, as only the identity hypothesis, but not the alternative model of P600 as a language-specific component, predict P3-to-P600 cross-decoding. Also, we would not interpret our results as indicating that *every* P600 can be *fully* accounted for by shared P3-patterns. Yet, the near-identical time-courses of within-P600 and P3-to-P600 decoding we observed demonstrate that P3-like patterns can account for the P600 effect to a degree that would be highly surprising under the non-identity hypotheses, but that is predicted by the identity hypothesis. This result calls direct domain-specific interpretations of P600 into question – particularly the still very frequent reverse inference concerning the presence of linguistic processes in non-linguistic experiments that elicit P600 effects (see Introduction section for recent examples).

Our conclusion that the P600 shares to a substantial degree neural processes with the P3 does by no means render late ERP positivities useless for studying language. However, it affects their role for informing neurocognitive models of language. It is generally established that the P3 marks linkage between salient stimuli and response selection (Nieuwenhuis et al., 2005). This interpretation should be considered when conceptualizing the neurocognitive processes underlying comprehension in complex domains such as language, music, or mathematics – in particular the processing of structural violations. Without any doubt, the cognitive processes so far attributed to the P600 – like syntactic structure building, reanalysis, or repair – *must* take place during language comprehension. However, the P600/P3, indicating overt registration of ungrammaticality or of words salient for other reasons, may be a precondition for – rather than an index of - initiating such second-order parsing processes (see Metzner, von der Malsburg, Vasishth, & Rösler, 2015, for compatible evidence). This can, for example, be used for investigations of individual differences in the sensitivity to linguistic contrast (e.g., at different stages of L2 acquisition; Tanner, McLaughlin, Hershensohn, & Osterhout, 2013; or see Tanner, 2018).

The P600-as-P3 account makes a range of testable predictions that would be inconsistent with the syntax-specific P600 model – in particular in domains where the syntax-specific account struggles. For example, as we discussed above, semantic manipulations induce a P600 when they are sufficiently salient (e.g., van de Meeredonk et al., 2011); this cannot be explained by a syntax-specific P600. We predict that these saliency-dependent semantic late positivities will show similar cross-decoding from P3 and syntactic P600s as we find here. However, other syntactic phenomena induce positive-going ERP components which are clearly not due to a single salient (i.e., P3-inducing) outlier event, such as those taxing syntactic working memory (e.g., Fiebach et al., 2002). Such positivities would *not* be expected to show cross-decoding with either a P300 or a violation-induced P600 of the extent we observed here. Thus, MVPA cross-decoding opens up a novel approach to examining several theoretical research questions in a quantitatively more explicit manner than was possible so far. In addition, this statistical technique might be useful for more specifically delineating the time courses of various cognitive resources relevant for sentence processing – including syntax, semantics, memory, but also (domain-unspecific) attention. In that vein, we have previously used time-resolved MVPA to demonstrate that the early (N400) and late (P600) components elicited by semantic violations in sentence contexts do not share patterns, but reflect entirely different processing stages (Heikel, Sassenhagen & Fiebach, 2018) – indicating a sequence of distinct, but partially overlapping processes underlying sentence processing.

In sum, we report that neural patterns corresponding to the P3 elicited by target detection in non-linguistic Oddball tasks re-occur during the P600 ERP component elicited by syntactic violations in sentences. This indicates substantial overlap of the neural patterns underlying P600 and P3, and strongly suggests that the P600 should be interpreted as a late, syntactic violation-dependent P3 (svP3). Neurocognitive models of language should accommodate this alternative interpretation of the P600. Observations of P600-like effects in other domains of perception should not automatically be interpreted as evidence for structural/language-related processing – calling into question straightforward reverse inference between domains. Our proposal stands up for empirical test. Proponents of a domain-specific interpretation of P600 should define falsifiable predictions that dissociate P600 and P3 at the neurocognitive level. Generalization-across-time decoding, as used in the present study, provides a valuable tool for further investigating such dissociations - e.g., to identify potential non-shared aspects of P600 and P3.

## Author Contributions

JS conceptualized the studies and analyzed data. JS and CJF wrote the manuscript.

## Acknowledgements

We thank Verena Willenbockel for providing stimulus material, Ingmar Brilmayer for discussion, Jean-Remí King for discussion and code, Joachim Scheel and Laura Nied for discussion and data collection, and Sebastien Nicolay and Cordula Hunt for data collection. We also thank three reviewers of a previous version of this manuscript for extensive constructive criticism. The research leading to these results has received funding from the European Community's Seventh Framework Programme (FP7/2013) under grant agreement n° 617891 awarded to CJF.

